# Trophic adaptation of large terrestrial omnivores to global change

**DOI:** 10.1101/2024.05.07.590891

**Authors:** Jörg Albrecht, Hervé Bocherens, Keith A. Hobson, Dorothée G. Drucker, Agnieszka Sergiel, Jon E. Swenson, Andreas Zedrosser, Adrian Marciszak, Elisabeth Iregren, Leena Drenzel, René Kyselý, Grzegorz Lipecki, Daniel Makowiecki, Jan Wagner, Tomasz Zwijacz-Kozica, Susanne A. Fritz, Eloy Revilla, Nuria Selva

## Abstract

Large omnivores at the top of food webs play a key role in ecosystems, as their ability to feed on multiple trophic levels stabilizes food-web dynamics and impacts ecosystem functioning. However, it is largely unexplored how large omnivores adapt their trophic interactions to altered resource availability under global change, particularly in terrestrial ecosystems. Here, we combine macroecological and paleoecological approaches and reveal that extant bears, the largest terrestrial omnivores, adapt their trophic position in food webs dynamically to net primary productivity and growing season length. Throughout their geographic ranges, extant bears occupy higher trophic positions in unproductive ecosystems with short growing seasons than in productive ecosystems with long growing seasons. Consistent with this geographic pattern, the trophic position of the brown bear sharply decreased at the transition from the Late Pleistocene to the Holocene, coinciding with an increase in net primary productivity and growing season length. These findings demonstrate that trophic interactions of omnivores are not static but change dynamically in response to environmental change. Our findings suggest that global change impacts on primary production and vegetation seasonality may trigger shifts in the functional role of omnivores, with consequences for food webs and ecosystem functions.

**Significance statement:** Large omnivores play a key role in ecosystems, as they stabilize food-web dynamics and impact ecosystem functioning. However, how omnivores adapt their trophic interactions to changes in resource availability under global change remains unexplored. Combining macroecological and paleoecological approaches, we show that extant bears, the largest terrestrial omnivores, occupy lower trophic positions in food webs as net primary productivity and growing season length increase. This trophic adaptation is evident across the geographic ranges of all extant terrestrial bear species and for brown bears at the transition from the Late Pleistocene to the Holocene. Therefore, impacts of global change on primary production and vegetation seasonality may trigger shifts in the functional role of omnivores, with consequences for food webs and ecosystems.

**One sentence summary:** Global changes in primary productivity and vegetation seasonality alter the functional role of large omnivores in terrestrial ecosystems.

## Introduction

Global change fundamentally reshapes the structure of terrestrial and aquatic food webs, which can have profound effects on entire ecosystems (1–3). Numerous studies have investigated the mechanisms behind changes in the structure of food webs under global change (1). However, an important mechanism that has received little attention is changes in trophic interactions due to the dynamic foraging behavior of consumers (4, 5). This mechanism is probably widespread and should be first detected in large omnivores at higher trophic levels, because they are adapted to rely on a wide range of resources, exhibit high behavioral flexibility, and often respond rapidly to environmental change (4, 5). As omnivores are common in food webs, a deterministic shift in the functional role of omnivores due to global change could result in the rewiring of entire food webs (4). In addition, changes in the functional role of omnivores, for instance from predation to herbivory, have direct consequences for food-web dynamics (5) and key ecosystem functions such as nutrient cycling, energy flux, and biomass production (6). Therefore, trophic responses of omnivores to global change could be a powerful indicator of critical transitions in food web structure and ecosystem functioning (4, 5).

Global change may exert particularly strong effects on the foraging behavior of omnivores via changes in resource availability. For instance, land-use intensification strongly reduces the availability of net primary productivity (NPP) to wildlife via the conversion of natural vegetation to livestock and crop production systems (7). On the other hand, anthropogenic nutrient inputs to ecosystems and food subsidies can increase the availability of resources to wildlife (8, 9). Moreover, prolonged growing seasons due to climate warming (10) can reduce seasonal bottlenecks in NPP. Food web theory suggests that the resulting changes in resource availability may cause shifts in the foraging behavior and trophic position of omnivores [‘dynamic omnivory’ hypothesis; Fig. 1; (4, 5, 11, 12)]. Yet, the trophic adaptation of large terrestrial omnivores to altered resource availability remains poorly understood (4, 5) because, traditionally, omnivory has been considered as a static trait (5) and studies that have investigated the dynamics of omnivory have mainly been conducted in micro- and mesocosm experiments or aquatic ecosystems (4, 5, 13, 14). In addition, most research has been conducted at local scales and over short periods, but little is known about the dynamics of omnivory at larger spatiotemporal scales (4, 5).

**Figure 1.**
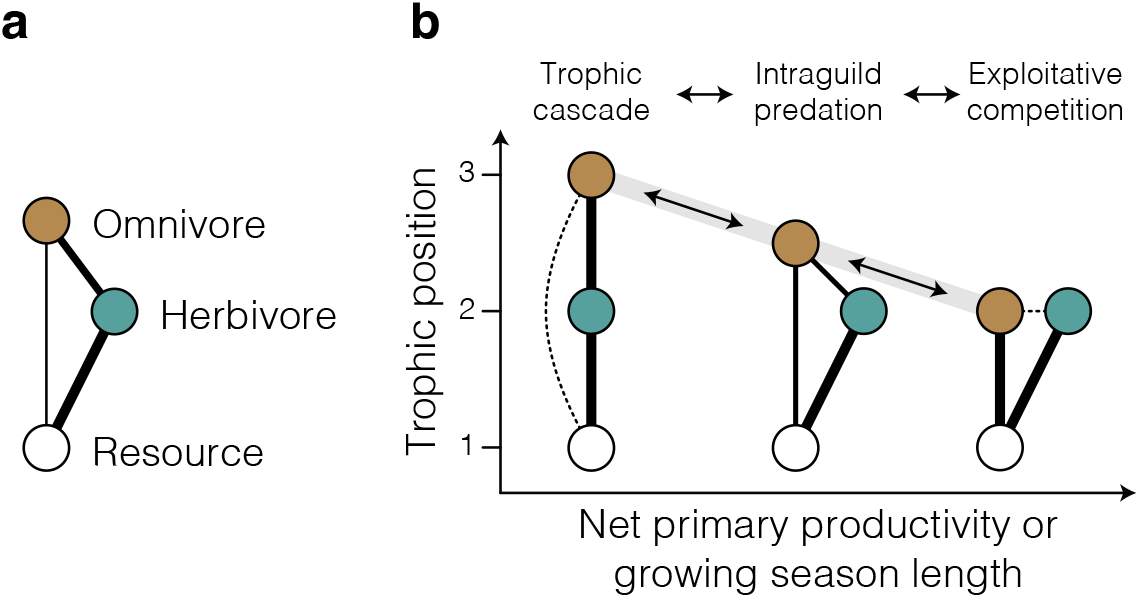
The trophic adaptation of omnivores to resource availability alters the structure of tritrophic food-web motifs. **a**, Schematic of a simple tritrophic food web with omnivory containing a basal resource, an herbivore and an omnivorous top predator. **b**, The ‘dynamic omnivory’ hypothesis from food web theory (5) suggests that with increasing resource availability at the base of food webs the structure of tritrophic food webs varies from a trophic cascade, where the top predator feeds exclusively on lower-level consumers, to intraguild predation, where the top predator feeds on both consumers and resources, to exploitative competition, where the top predator feeds exclusively on the resource and competes with the lower-level consumer for this resource. Dashed lines in **b** depict indirect interactions.

Here, we aim to fill these knowledge gaps by combining macroecological and paleoecological approaches to investigate how large terrestrial omnivores adapt their trophic position in food webs to changing NPP and growing season length at global and millennial scales. We leverage the unique insights gained by both approaches (15, 16) to infer how large terrestrial omnivores adapt their trophic position in food webs to global change. We focus on the seven extant terrestrial bear species (Order: Carnivora, Family: Ursidae), which are the largest terrestrial omnivores and occupy a wide range of biomes from the arctic tundra to tropical rainforests. Unlike most other large carnivores, bears show a preference for low-protein diets and have relatively weak craniodental adaptations to carnivory (17, 18), which allows them to maintain a high degree of dietary flexibility. Owing to their broad dietary niches, bears contribute to a multitude of ecosystem processes, such as predation, scavenging, or frugivory that can have strong impacts on prey populations (19), plant regeneration (20, 21), nutrient cycling (22, 23) and energy fluxes (6) within and across terrestrial and aquatic ecosystems.

## Results

We first investigated how the trophic position of extant bears is related to NPP and growing season length across their geographic ranges (Fig. 2). To do so, we compiled a comprehensive database of dietary compositions based on micro-histological analyses of fecal and stomach contents throughout the geographic ranges of the seven extant terrestrial bear species, using 210 records from 155 studies (Fig. 2a) (24). We excluded the polar bear (*U. maritimus*) from our analysis, because this species almost exclusively hunts for marine prey on Arctic Sea ice, but fasts on land during the ice-free season (25). Therefore, the minor contribution of terrestrial food sources during the ice-free season is not representative of the species’ trophic niche (26–28). Based on the dietary data for the remaining seven terrestrial bear species, we used a Bayesian hierarchical model to estimate the trophic position as the percentage of the dietary energy that is contributed by animal prey (including vertebrates and invertebrates), while accounting for the effects of digestibility and energy content of different food sources (Fig. 2b; *Materials and Methods*; *SI Appendix*, Table S1). In a second step, the model related the estimated trophic position to NPP (kg C m^−2^ a^−1^) and meteorological growing season length (29) (months with mean temperature *T* > 0°C; hereafter growing season length) at each of the study locations (Fig. 2c,d). The temperature threshold of 0°C demarcates the growth period of frost-resistant plants (30), as well as the active period of bear populations in temperate and boreal biomes [e.g., *Ursus arctos* and *U. americanus* (31, 32)]. To account for the potential effects of competition between sympatric bear species, we also included a factor indicating whether a focal population was located within the range of another larger (dominant) or smaller (subordinate) bear species (33, 34). The model included random factors for study and species identity to account for the non-independence of data from the same species and study and propagated all uncertainties associated with the estimated trophic position through the entire model.

**Figure 2.**
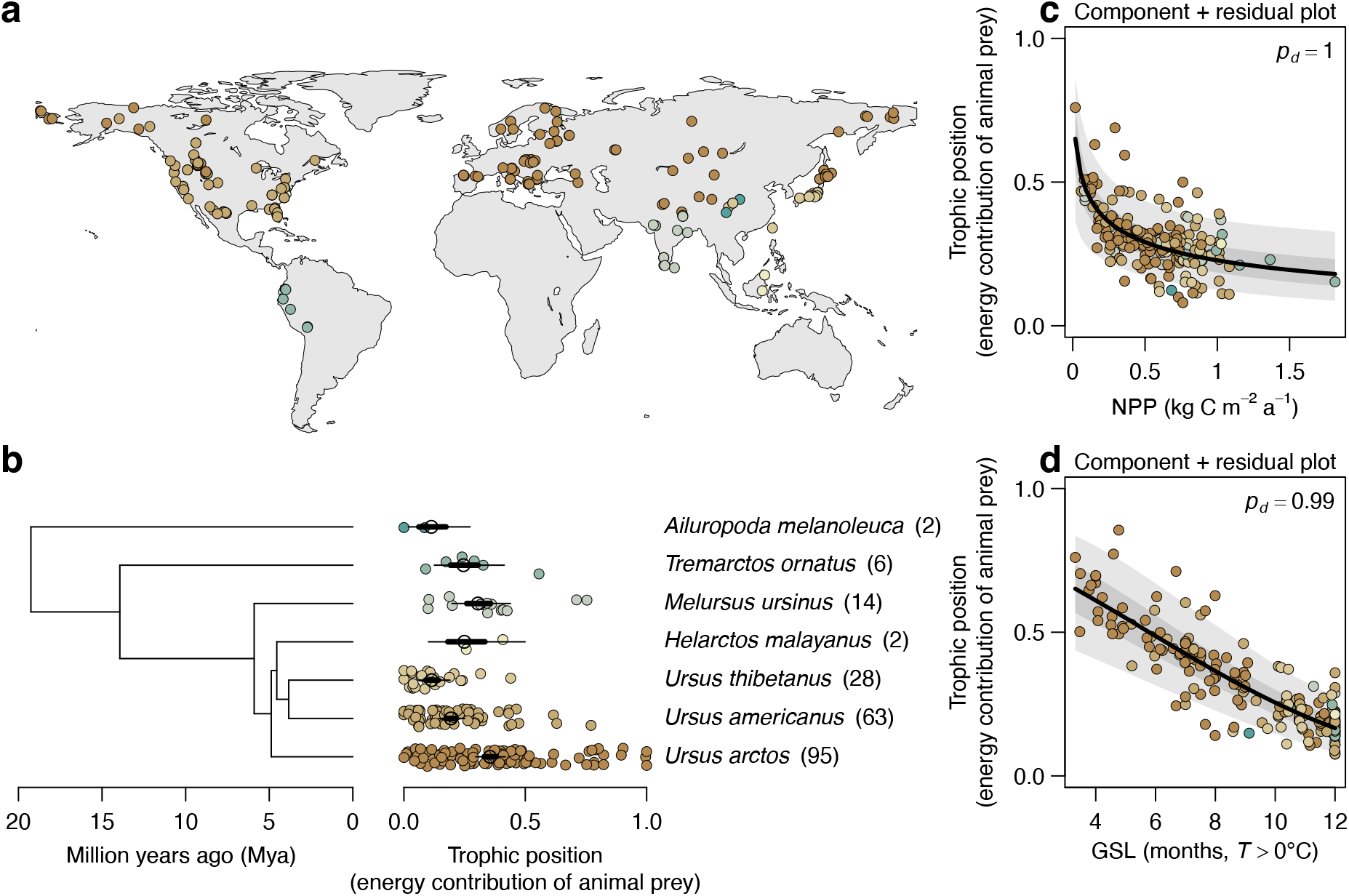
Global variation in the trophic position of bears. **a**, Geographic distribution of diet studies based on micro histological analyses of faecal and stomach content included in the meta-analysis. **b**, Phylogenetic tree of the lineage of bears (excluding *U. maritimus*) (76) alongside species-specific trophic position, estimated as the relative dietary energy contribution of animal prey. The phylogenetic tree is plotted for illustrative purposes only. Black circles, thick, and thin lines represent posterior estimates (median, 50% and 90% ETIs [Equal-Tailed Intervals]) of species-specific trophic position, whereas small circles represent estimated trophic position in each study location. Numbers in brackets give the sample size for each species. **c, d** Component + residual plots showing the partial relationships of trophic position with (**c**) net primary productivity (NPP, kg C m^−2^ a^−1^) and (**d**) meteorological growing season length (GSL, months with *T* > 0°C) while conditioning on the effects of the other predictor. Black lines, dark and light gray bands represent estimated relationships and uncertainty (median, 50% and 90% ETIs, respectively). Sample sizes are *n*_observations_ = 210, *n*_study_ = 155, *n*_species_ = 7. *p*_*d*_ is the posterior probability of a relationship being the same direction as the median.

The model revealed that the trophic position of extant bears across their geographic ranges is negatively related to NPP (*p*_*d*_ = 1, posterior probability of the relationship being negative) and growing season length (*p*_*d*_ = 0.99; Fig. 2c,d; *SI Appendix*, Table S2). Thus, terrestrial bears generally occupy higher trophic positions in ecosystems with low productivity and short growing seasons and lower trophic positions in ecosystems with high productivity and long growing seasons. A decline in trophic position may be attributable to (i) a population effect, where local bear populations alter their foraging behavior relative to NPP and growing season length, or (ii) a species occurrence effect, where bear species with a certain trophic position simply do not occur in environments with a particular NPP and growing season length. To disentangle these two effects, we ran an additional model where we separated the effects of NPP and growing season length into these two components (*Materials and Methods*). The two-component model showed that variation in trophic position was exclusively explained by population effects, and not by the species occurrence effect (*SI Appendix*, Table S2). These effects were robust despite additional effects of competition with sympatric bear species, which caused a decrease in the trophic position of subordinate bear species (*p*_*d*_ = 0.99; *SI Appendix*, Table S2). This indicates that the trophic position of extant bears is a flexible trait that emerges from adaptations to local resource availability and interspecific competition with other sympatric bear species at the population level.

In a second step, we investigated how European brown bears adapted their trophic position to the marked increases in NPP and growing season length at the transition from the Late Pleistocene to the Holocene (Fig. 3a). To do so, we used a comprehensive collection of fossil and subfossil bone and tooth remains of brown bears (*n* = 219) and red deer (*Cervus elaphus, n* = 372) across Europe, covering the last 55 ka BP (Fig. 3b) (24). We estimated the trophic position of brown bears based on δ^15^N (‰) of collagen through time using a Bayesian hierarchical model, in which the δ^15^N values of red deer from the same region were included as a dietary baseline for a strict herbivore (Fig. 3c; *Materials and Methods*). The model accounted for the effects of altitude (35) and the type of sampled material [bones or teeth (34)] on δ^15^N values of brown bears and red deer and propagated all uncertainties associated with these factors, as well as with trophic fractionation (34, 36) through the entire model (*Materials and Methods*; *SI Appendix*, Table S3). In the last step, the model estimated the relationship of the trophic position of brown bears with NPP and growing season length, which were derived from global climate and vegetation models [HadCM3 and BIOME4 (37)] (Fig. 3a).

**Figure 3.**
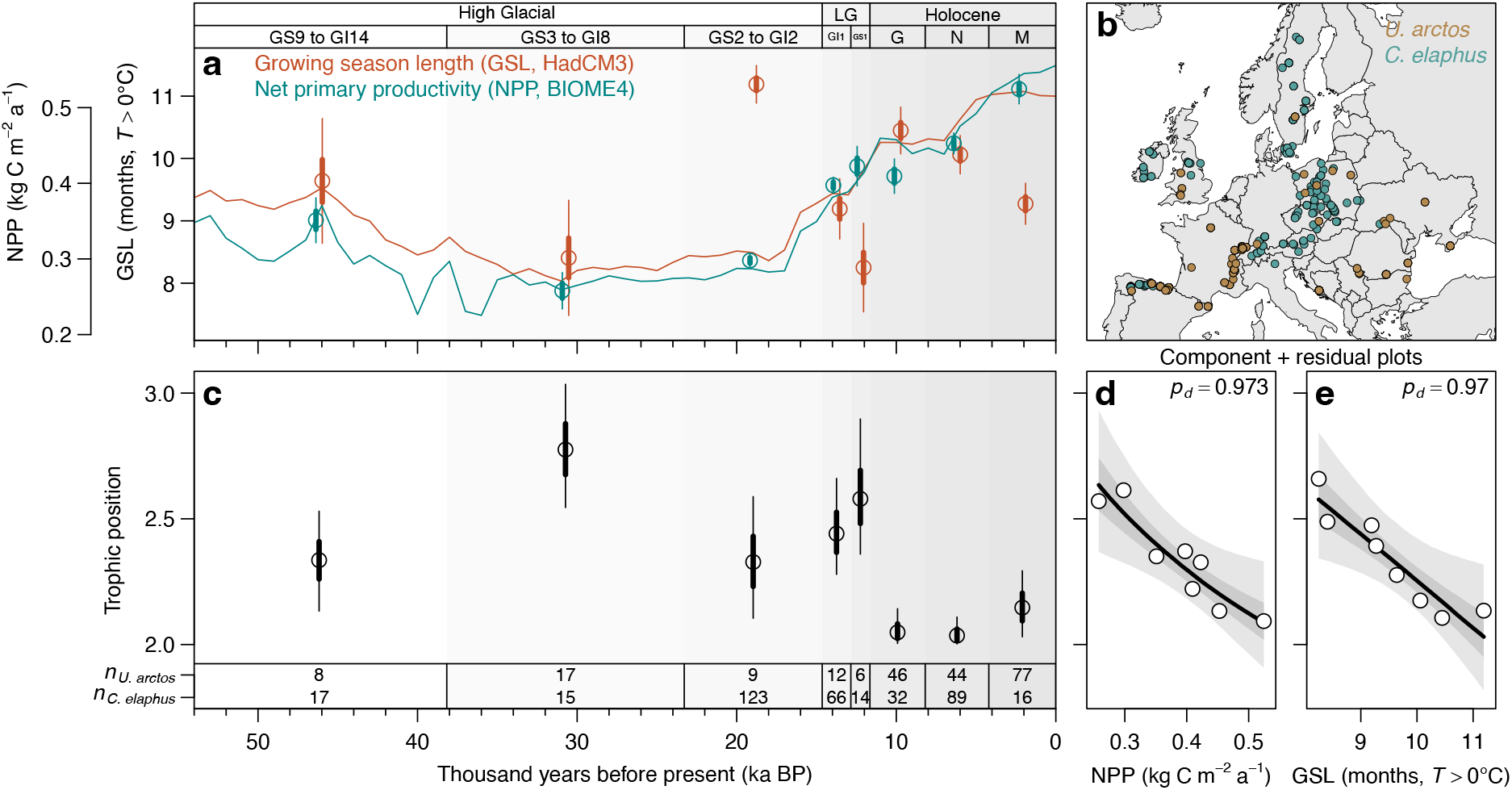
Change in trophic position of the brown bear (*Ursus arctos*) from Late Pleistocene to Holocene. **a**, Net primary productivity (NPP, kg C m^−2^ a^−1^) and meteorological growing season length (GSL, months with *T* > 0°C), based on global climate and vegetation models [HadCM3 and BIOME4 (37)] during the last 55,000 years. Circles, thick and thin error bars represent mean NPP and growing season length, as well as 50% and 90% CIs [Confidence Intervals], across the subfossil bone and tooth samples of brown bear and red deer (*Cervus elaphus*) in each time bin (*Materials and Methods*). Lines indicate mean temporal trends in NPP and growing season length across all sample locations shown in (**b**). Abbreviations are LG, Late Glacial; GS9 to GI14, Greenland Stadial 9 to Greenland Interstadial 14; GS3 to GI8, Greenland Stadial 3 to Greenland Interstadial 8; GS2 to GI2, Greenland Stadial 2 to Greenland Interstadial 2; GI1, Greenland Interstadial 1, Greenland Stadial 1; M, Meghalayan; N, Northgrippian; G, Greenlandian. **b**, Locations of subfossil bone and tooth samples of brown bear and red deer across the European continent. **c**, Trophic position of the brown bear as estimated by a Bayesian hierarchical model with red deer as baseline, where a value of 2 corresponds to a strict herbivore (i.e., trophic level = 2) and values larger than 2 reflect an increasing dietary contribution of animal prey. Black circles, thick and thin lines represent posterior estimates (median, 50% and 90% ETIs, respectively). Sample size (*n*) in each time bin is given at the bottom of the graph. **d**,**e** Component + residual plots showing the partial relationships of trophic position with NPP and growing season length, conditioned on the effect of the other predictor variable. Black lines, dark and light gray bands represent estimated relationships and uncertainty (median, 50% and 90% ETIs, respectively). *p*_*d*_ is the posterior probability of a relationship being the same direction as the median.

We found that European brown bears occupied higher trophic positions during the Late Pleistocene than during the Holocene (Fig. 3c). This observed decrease in trophic position of European brown bears from the Late Pleistocene to the Holocene was closely associated with increasing NPP (*p*_*d*_ = 0.97) and growing season length (*p*_*d*_ = 0.97) at the onset of the Holocene (Fig. 3d,e; *SI Appendix*, Table S3). Consequently, the occupied trophic positions were highest during the period characterized by the lowest NPP and shortest growing seasons (period GS3 to GI8 in Fig. 3c) and lowest during those periods characterized by high NPP and long growing seasons (periods Greenlandian and Northgrippian in Fig. 3c). Most strikingly, the paleoecological patterns for brown bears closely resembled the macroecological patterns observed across the geographic ranges of the seven extant terrestrial bear species (Figs. 2 and 3).

## Discussion

The observed trophic adaptation to NPP and growing season length is in line with predictions from food web models and with empirical observations from micro- and mesocosm experiments and lake ecosystems (5, 11–14). We attribute the observed trophic adaptation to two nonexclusive mechanisms: (i) tracking of seasonal peaks in plant productivity (e.g., fruit and seed production), which is facilitated by the mobility of these large omnivores (4, 5); and (ii) shifts in foraging preferences if foraging for plant resources is energetically more efficient than foraging for animal prey, due to low search and handling times in dense vegetation patches (38, 23). This is supported by previous work reporting rapid trophic adaptations within some brown bear populations to seasonal and annual variability in resource availability (23, 39). Our study highlights that, similar to aquatic food webs, the trophic interactions of large omnivores in terrestrial food webs are not static but can change dynamically in response to altered resource availability. The fact that extant terrestrial bears show a consistent trophic response to resource availability across their geographic ranges puts the findings of these previous studies into a macroecological perspective. Over long time scales, the observed trophic adaptation to resource availability throughout the geographic ranges of bears might even be a driver of intraspecific genetic differentiation and population structure (40, 41).

The sharp decrease in the trophic position of European brown bears at the onset of the Holocene (11.7 ka BP) challenges the hypothesis that the trophic position of the European brown bear decreased due to competitive release after the extinction of the cave bear [*Ursus spelaeus*; 24.3–26.1 ka BP (34, 42)]. Our results rather suggest that European brown bears primarily adapted their trophic niche to enhanced resource availability, whereas competitive release seemingly played a secondary role. This is supported by the fact that we found consistent effects of NPP and growing season length across extant terrestrial bears after accounting for the potential effects of competition between sympatric species (*SI Appendix*, Table S2). The relatively steady trophic position of European brown bears throughout the Early- and Mid-Holocene highlights that trophic adaptations can develop into stable trophic strategies that persist within populations when environmental conditions remain constant (Fig. 3c). Overall, the strong agreement between the macroecological and paleoecological approaches suggests that the trophic position occupied by extant bears in food webs across their current geographic ranges is the result of trophic adaptations to local resource availability.

The observed trophic adaptation of brown bears to environmental change at the onset of the Holocene (Fig. 3) indicates that current global change may have profound effects on the trophic position and functional role of omnivores in terrestrial food webs. For instance, land-use intensification reduces the availability of primary production to wildlife (7) and may cause opportunistic shifts in foraging behavior to alternative food sources, including livestock or crops (43). In addition, anthropogenic nutrient inputs to ecosystems and food subsidies increase the availability of resources, which can also affect the foraging behavior and trophic interactions of omnivores (9). Finally, the increasing length of growing seasons associated with climate warming (10) may release omnivores from seasonal bottlenecks in primary production and may cause dietary shifts from animal prey to plant resources (5, 44). The release from seasonal bottlenecks in NPP associated with warmer autumn and spring temperatures also affects the activity and wintering behavior of bears (31, 32), which might in part explain shortened hibernation periods and cases of non-hibernation in some bear populations (45). Despite the signal of trophic adaptation of populations to environmental change in our study, it remains unclear to what extent populations of species that live at the physiological or ecological limits of their range in extreme environments (e.g., arctic, desert, or alpine environments) are able to cope with future changes in environmental conditions. Their adaptive capacity will depend on the magnitude and velocity of environmental change, as well as on whether alternative food sources meet the nutritional requirements of consumers (4, 46). For instance, it has been suggested that the switch of polar bears from marine prey to terrestrial food sources (e.g., berries or bird nests) in response to reduced sea-ice extent due to climate change is associated with declines in body size, body condition and reproduction (47).

A growing body of literature highlights the pervasive impact of large animals on the structure and functioning of Pleistocene ecosystems and the cascading effects of historic declines and losses of this megafauna on ecosystems (48). Our study adds another dimension to these studies because the observed trophic adaptation of large terrestrial omnivores to altered resource availability under global change could have impacts on food webs and ecosystem functions that go beyond the effects of population declines. On the one hand, food-web models indicate that rapid responses of omnivores to changing resource availability stabilize population dynamics under global change (5). Despite these potentially stabilizing effects, changes in the trophic position of large omnivores directly affect the structure of food webs (e.g., food chain length and interaction strengths) and have strong effects on nutrient cycling, energy fluxes, and biomass stocks in ecosystems (6). Apart from these direct impacts on food webs and ecosystem functions, the trophic adaptation of omnivores may also translate into a higher frequency of human–wildlife conflicts, if omnivores increasingly use anthropogenic resources in agricultural landscapes (43).

## Conclusion

By combining macroecological and paleoecological approaches, our study provides strong empirical evidence that large terrestrial omnivores adapt their trophic position in food webs dynamically to resource availability, with a general decrease in trophic position in response to higher primary production and longer growing seasons. Our findings complement empirical observations from previous small-scale experiments and aquatic ecosystems, suggesting that the trophic adaptation of omnivores to resource availability is common across aquatic and terrestrial ecosystems. Because the trophic position of omnivores in food webs is closely linked to their functional role in ecosystems, their trophic adaptation to current global change is likely to have cascading effects on the structure of food webs and ecosystem functions.

## Materials and Methods

### Macroecological analysis

#### Trophic data

To obtain dietary data for the seven extant terrestrial bear species, we conducted a literature search in the Web of Science Core Collection for publications until 2018. We searched titles and abstracts using the keywords “(food *OR* feed* *OR* diet* *OR* forag* *OR* nutri* *OR* scat* *OR* fec* OR faec* *OR* stomach* *OR* gut*) *AND* (ursidae *OR* ursus *OR* melursus *OR* tremarctos *OR* helarctos *OR* ailuropoda *OR* bear* *OR* panda)”. We also checked the reference lists of the retrieved publications for additional studies that had not been identified by the keyword search. To be included in our analysis, the studies had to meet the following criteria: (i) Diet assessment was based on microhistological analyses of food remains in scats or stomachs. (ii) Sample collection in the field covered the entire seasonal activity period of the focal species. (iii) All food items, instead of only a subset, were reported. (iv) Reported food items could be mapped onto established macroecological diet classification schemes (49). (v) The number of collected samples was reported. (vi) Sufficient data to calculate one of the following measures was reported: relative frequency of occurrence of food items (*F*_*i*_), calculated as the number of occurrences of food item *i* divided by the total number of occurrences of all food items 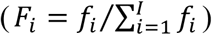; the relative volume of food items (*V*_*i*_), calculated as the mean volume of food item *i* in scats or stomachs divided by the total volume of all food items, that is 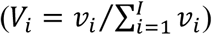; the relative dietary contribution based on ingested dry weight of food items (*D*_*i*_), calculated as the product of the relative volume of food item *i* and correction factors *c*_*Di*_ that account for differences in the digestibility of food items and normalizing to proportions (*D*_*i*_ = *c*_*Di*_*V*_*i*_/∑_*i*_ *c*_*Di*_*V*_*i*_); or relative dietary energy contribution of food items (*E*_*i*_), calculated as the product of the relative dietary dry weight contribution of food item *i* and correction factors *c*_*Ei*_ that account for differences in the energy content of food items and normalizing to proportions (*E*_*i*_ = *c*_*Ei*_*Di*/∑*i c*_*Ei*_*D*_*i*_). We excluded the polar bear from our analysis, because this species almost exclusively hunts for marine prey on Arctic Sea ice, but fasts on land during the ice-free season (25). Therefore, the minor contribution of terrestrial food sources during the ice-free season is not representative of the species’ trophic niche (26–28).

In total, 155 publications met the above criteria, including articles in scientific journals, master’s and PhD theses, and gray literature (e.g., non-commercial publications in the form of technical reports or conference proceedings) (24). For each publication, we extracted the geographic coordinates of the study location, the type of collected samples (i.e., scats or the stomachs of dead individuals), the number of samples, and the reported food items in scats or stomachs. We mapped the reported food items onto one of 20 dietary categories based on an established macroecological diet classification scheme (49) (*SI Appendix*, Table S1) and recorded the quantity of food items in the diet using the above described variables. To convert estimates of relative volume into relative dietary contribution or dietary energy contribution, we compiled correction factors that account for differences in the digestibility and energy content of food items from the literature (*SI Appendix*, Table S1).

#### Environmental variables

For the geographic location of each record, we extracted data on net primary productivity (NPP, kg C m^−2^ a^−1^) [Terra MODIS17A3, spatial resolution: 0.5° (50)] and near-surface daily average air temperature at monthly resolution [CHELSA, spatial resolution: 0.5° (51)]. We calculated growing season length as the number of months, in which the mean temperature was above 0°C. This temperature threshold demarcates the thermal growth period of frost-resistant plants (30), as well as the active period of bear populations in temperate and boreal biomes [e.g., *Ursus arctos* and *U. americanus*, respectively (31, 32)].

#### Macroecological model

To analyze global patterns of trophic position across the seven extant terrestrial bear species, we used a Bayesian hierarchical model that consisted of two sub-models. Sub-model I was used to (i) represent the relative volume of food items in sampled materials for those datasets with missing information on relative volume (107 of 210 datasets), based on the close relationship with the relative frequency of occurrence of food items (*SI Appendix*, Fig. S1) and (ii) to estimate the relative dietary energy contribution of animal prey, including vertebrates and invertebrates (i.e., the trophic position). All uncertainties associated with the data imputation and estimation of trophic position were propagated through the entire model. Sub-model II was then used to estimate the relationship of the trophic position with NPP and growing season length. To inform sub-model I, we first quantified the relationship between the relative frequency of occurrence (*F*) and relative volume (*V*) for those datasets (i.e., unique study-year or study-location combinations), in which both variables had been reported (*n* = 106 datasets). For each dataset *j*, we quantified the relationship between *F* and *V* using geometric mean regression to account for the fact that both variables are subject to measurement error (52). We calculated the geometric mean slope as 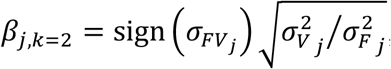, and the associated intercept as 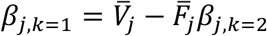, where 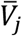 and 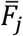 are the mean values of both variables for dataset *j*. The distribution of geometric mean slopes and associated intercepts across the datasets was then used to inform the data imputation in sub-model I. In particular, sub-model I assumed that the slopes and intercepts follow a multivariate normal distribution around 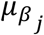 and variance-covariance matrix Σ_*β*_:

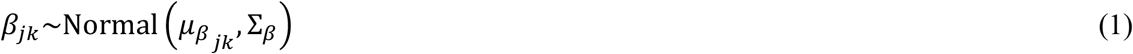

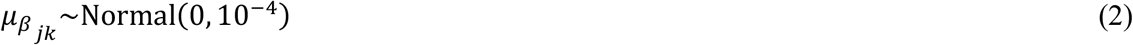

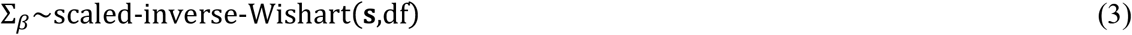

where Σ_*β*_ is a 2 × 2 variance–covariance matrix. We used a noninformative scaled inverse Wishart prior (53) with scale *s*_*k*_ = 1 and d.f. = 2. This prior follows a half-*t* distribution for the standard deviation, *σ*_*kk*_ ∼ *st*^+^_d.f._, and has a marginal uniform prior distribution for the correlation parameter *ρ*_*kl*_. For each dataset *j* without information on *V*_*i*[*j*]_ the model then drew an intercept and slope from the multivariate distribution and estimated values of *V*_*i*[*j*]_ using the observed values of *F*_*i*[*j*]_ based on the following regression formula:

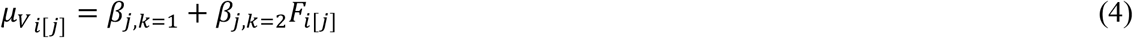

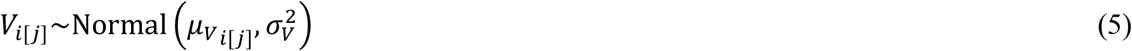

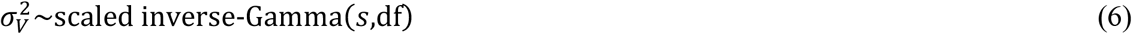

We modeled *V*_*i*[*j*]_ using a normal distribution with mean 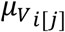 and residual variance 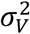. For the residual variance, we used a weakly informative scaled inverse gamma prior (54) with scale *s* = 1 and two degrees of freedom (d.f. = 2), which for the standard deviation follows a half-*t* distribution. Then the relative dietary energy contribution *E* of each food item was calculated for each dataset *j* by multiplying the values of *V* with correction factors *c*_*D*_ *and c*_*E*_ that account for differences in the digestibility and energy content of food items and normalizing to proportions:

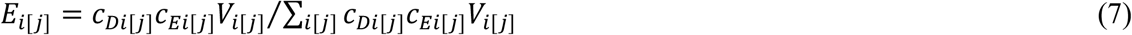

Based on the relative dietary energy contribution *E* of each food item, we calculated for each dataset *j* the trophic position as the relative dietary energy contribution of animal prey *P*_*j*_ (including vertebrates and invertebrates). The process model then quantified the relationship of *P* with NPP and growing season length. The model can be written as follows:

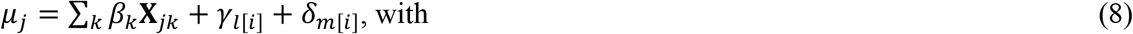

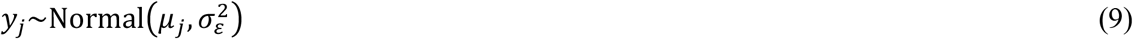

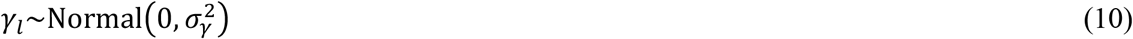

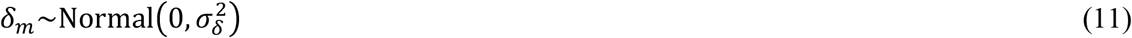

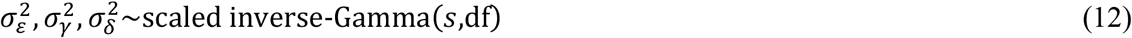

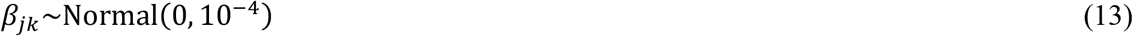

where *μ*_*j*_ is the expected value of the *j*th observation, **X** is a model matrix including an intercept and the two explanatory variables (NPP [log-transformed] and growing season length) and an associated vector of parameters *β*, and *γ*_*l*_ is a random effect for the *l*th study, which is normally distributed around 0 with variance *σ*_*γ*_^2^, and *δ*_*m*_ is a random effect for the *m*th species, which is normally distributed around 0 with variance *σ*_*δ*_^2^. We modeled the trophic position with a normal distribution around *μ*_*j*_ and residual variance *σ*_ε_^2^. For modeling purposes, we first compressed values of *P*_*j*_ which were in the closed interval (0 ≤ *y* ≤ 1) into the open interval (0 < *y* < 1) and subsequently applied a probit transformation: *y*_*j*_ = probit((*P*_*j*_(*n* – 1) + 0.5)/*n*), in which *n* is the total number of observations (55). The compression was necessary, because the probit function is defined on the open interval (0 < *y* < 1; range of *P*_*j*_ after transformation: 0.00034–0.99934). We used an uninformative normal prior with a mean of 0 and a variance of 10^4^ for the fixed effects and weakly informative scaled inverse gamma priors with scale *s* = 1 and two degrees of freedom (d.f. = 2) for variances of random effects and residuals.

### Paleoecological analysis

#### Database of subfossil and fossil remains

To reconstruct the trophic position of the European brown bear during the Late Pleistocene and Holocene, we compiled a database of 591 dated and georeferenced subfossil and fossil remains of brown bears and red deer using published sources (*n* = 483 samples) and new material (*n* = 108) (24). Dates of specimens were based on direct radiocarbon (^14^C) dating (*n* = 237) or based on the context of the excavation sites (*n* = 354). Contextual dates were either obtained based on related artifacts (*n* = 196) or in case of three sites (El Miron, Kiputz IX, and Emine-Bair-Khosar) based on age-depth models (*n* = 158). Radiocarbon dating of new fossil material (*n* = 55) was done using accelerator mass spectrometry at the Curt Engelhorn Centre for Archaeometry (CEZA), Mannheim, Germany. Dated fossils without geolocations were geocoded manually using the name of the fossil site. The quality and reliability of all radiocarbon dates was assessed based on dating method, stratigraphy, association and material (56). This resulted in 219 samples of brown bear and 372 samples of red deer. Age-depth models were fitted using package *Bchron* [version 4.7.6 (57)] in *R* [version 4.0.3 (58)]. The ^14^C ages of these fossils were calibrated using package *Bchron* and the IntCal13 curve (59).

#### Stable isotope analysis

Collagen was extracted from subfossil and fossil specimens following previously established protocols (60, 61). The extraction process included a step of soaking in 0.125 M NaOH between the demineralization and solubilization steps to achieve the elimination of lipids and humic acids. Elemental analysis (C_coll_, N_coll_) and isotopic analysis (δ^13^C_coll_, δ^15^N_coll_) were conducted at two laboratories. At the Department of Geosciences of Tübingen University, Germany, an NC2500 CHN-elemental analyzer was coupled to a Thermo Quest Delta+ XL mass spectrometer. At the Environment Canada Stable Isotope Laboratory in Saskatoon, Canada, collagen was combusted at 1030°C in a Carlo Erba NA1500 or Eurovector 3000 elemental analyser coupled with an Elementar Isoprime or a Nu Instruments Horizon isotope ratio mass spectrometer. The international standard for δ^13^C measurements was the Vienna Pee Dee Belemnite (VPDB) carbonate that for δ^15^N atmospheric nitrogen (AIR). Analytical error, based on within-run replicate measurement of laboratory standards (Saskatoon: BWBIII keratin and PRCgel; Tubigen: albumen, modern collagen, USGS 24, IAEA 305 A), was ±0.1‰ for δ^13^C values and ±0.2‰ for δ^15^N values. Reliability of the δ^13^C_coll_ and δ^15^N_coll_ values was established by measuring its chemical composition, with C/N_coll_ atomic ratio ranging from 2.9 to 3.6 (62), and percentage of C_coll_ and N_coll_ above 8% and 3% (63), respectively.

#### Trophic discrimination factors and corrections for material type and elevation

We obtained (i) data on trophic discrimination factors based on bone collagen of predators and their prey, (ii) reference data to model the relationship between elevation and the isotope composition of plants and herbivores, as well as (iii) data on differences in the isotopic composition of collagen from bone and teeth from the literature (34–36, 24).

#### Environmental variables

We used package *pastclim* [version 2.0 (64)] to annotate each sample with spatiotemporally explicit data on net primary productivity (NPP, kg C m^−2^ a^−1^) from a global dynamic vegetation model [BIOME4, spatial resolution: 0.5° (37)], near-surface daily average air temperature at monthly resolution from a global climate model [HadCM3, spatial resolution: 0.5° (37)], and with data on elevation from a global elevation model [GMTED 2010, spatial resolution: 0.0083° (65)]. Analogous to the macroecological analysis, we calculated growing season length as the number of months in which the near-surface daily average air temperature was above 0°C.

#### Temporal resolution of analysis

For the analysis, we defined eight time periods that were used to bin the data (Fig. 3): Meghalayan (0–4.2 ka BP), Northgrippian (4.2–8.2 ka BP), Greenlandian (8.2–11.7 ka BP), Greenland Stadial 1 (GS-1, 11.7–12.8 ka BP), Greenland Interstadial 1 (GI-1, 12.8–14.6 ka BP), Greenland Stadial 2 to Greenland Interstadial 2 (GS-2 to GI-2, 14.6–23.3 ka BP), Greenland Stadial 3 to Greenland Interstadial 8 (GS-3 to GI-8, 23.3– 38.2 ka BP), Greenland Stadial 9 to Greenland Interstadial 14 (GS-9 to GI-14, 43.8–54.2 ka BP). These time bins represent a trade-off between temporal resolution and sample size within each time period. The Subdivision of Late Pleistocene and Holocene time periods was based on current definitions of stratigraphic stages (66, 67).

#### Paleoecological model

To analyze changes in the trophic position of the brown bear during the past 55,000 years, we used a Bayesian hierarchical model that consisted of two sub-models. Sub-model I was used to estimate the trophic position of the brown bear within each time period using the δ^15^N values of red deer collagen as a dietary baseline for a strict herbivore. The model accounted for the effects of altitude (35), the type of sampled material [bones or teeth (34)] on δ^15^N values of brown bears and red deer, and propagated all uncertainties associated with these factors as well as with trophic fractionation (34, 36) through the entire model. Sub-model II then related the estimated trophic position to NPP and growing season length. Below we describe the model in detail.

First, sub-model I estimated the average difference between the isotopic value of tooth and bone material based on paired samples of bone and teeth from the same individuals of different species of large carnivores [n = 35 (24, 34)]:

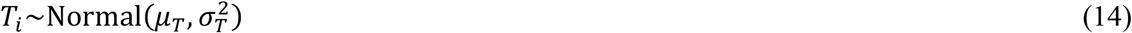

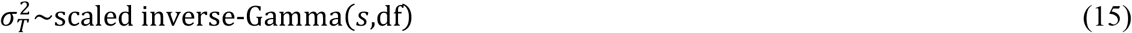

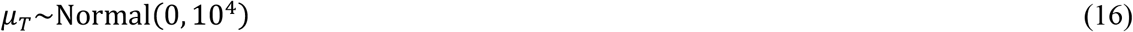

We modeled the observed difference between the isotopic values of tooth and bone material *T*_*i*_ using a normal distribution with mean *μ*_*T*_ and residual variance 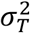. We used an uninformative normal prior with a mean of 0 and a variance of 10^4^ for *μ*_*T*_ and a weakly informative scaled inverse gamma prior with scale *s* = 1 and two degrees of freedom (d.f. = 2) for 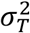.

Second, sub-model I estimated the effect of elevation on the isotopic values based on data that relates δ^15^N (‰) of plants and herbivorous animals to elevation [n = 69 (24, 35)]:

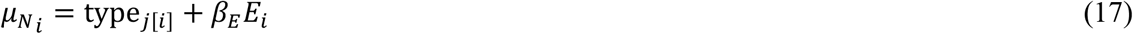

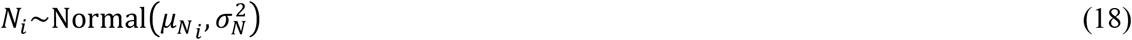

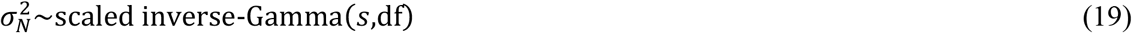

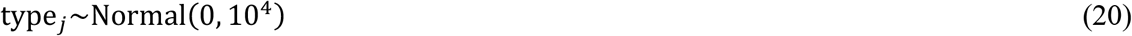

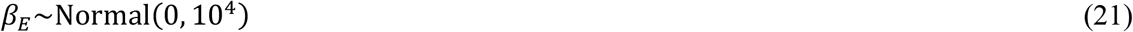

where 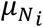 is the expected δ^15^N value of the *i*th observation, type_*j*[*i*]_ is the type of material (vegetation, sheep wool, cattle hair, or goat hair), *β*_*E*_ is the effect of elevation, and *E*_*i*_ is the elevation at which sample *i* has been collected. We modeled the observed δ^15^N values *N*_*i*_ using a normal distribution with mean 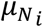 and variance 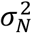. We used uninformative normal priors with a mean of 0 and a variance of 10^4^ for the effects of material type and elevation and a weakly informative scaled inverse gamma prior with scale *s* = 1 and two degrees of freedom (d.f. = 2) for 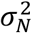.

Third, sub-model I used the estimates of *β*_*E*_ and *μ*_*T*_ to estimate the mean bias-corrected δ^15^N values for red deer (baseline) and brown bear (consumer) in each of the eight time periods (*SI Appendix*, Fig. S2):

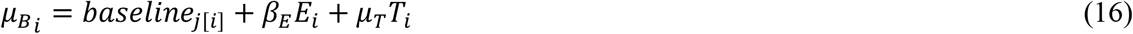

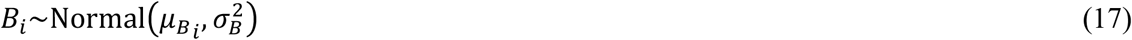

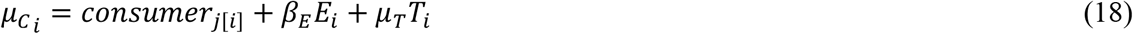

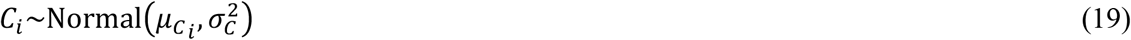

where 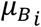 and 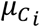 are the expected δ^15^N values of red deer and brown bear samples, baseline_*j*_ and consumer_*j*_ are the mean bias-corrected nitrogen signatures of red deer and brown bear in time period *j*, respectively, *E*_*i*_ is the elevation at which sample *i* has been collected, and *T*_*i*_ indicates whether the sample is a tooth (*T*_*i*_ = 1). We modeled the observed δ^15^N values of red deer *B*_*i*_ and brown bear *C*_*i*_ using normal distributions with means 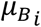 and 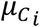 and variances 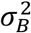 and 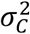, respectively. We used uninformative normal priors with a mean of 0‰ and a variance of 10^4^‰ for the mean bias-corrected δ^15^N values (baseline_*j*_ and consumer_*j*_) and weakly informative scaled inverse gamma priors with scale *s* = 1 and two degrees of freedom (d.f. = 2) for the variances (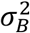 and 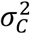):

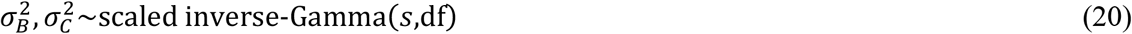

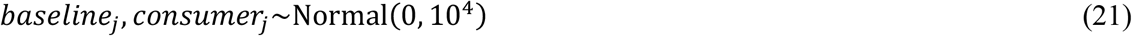

The trophic position of the brown bear in period *j TP*_*j*_ was then estimated as the difference between the mean bias-corrected δ^15^N values of brown bear and red deer, divided by the trophic discrimination factor *μ*_Δ_ plus an offset for the trophic level of the baseline [*λ* = 2 (68)]:

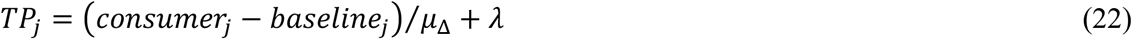

The trophic discrimination factor was estimated based on data for pairs of large carnivores and their prey [n = 10 pairs (24, 34)]:

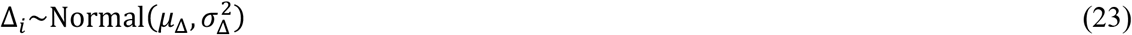

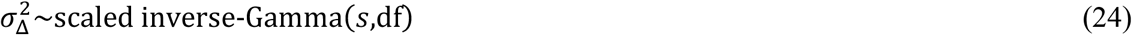

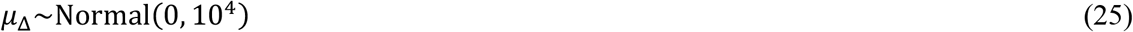

We modeled the observed trophic discrimination factor Δ_*i*_ using a normal distribution with mean *μ*_Δ_ and residual variance 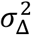. We used an uninformative normal prior with a mean of 0 and a variance of 10^4^ for *μ*_Δ_ and a weakly informative-scaled inverse gamma prior with scale *s* = 1 and two degrees of freedom (d.f. = 2) for 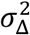.

In a final step, sub-model II related the estimated trophic position *TP*_*j*_ in each period *j* to the NPP and growing season length in each period:

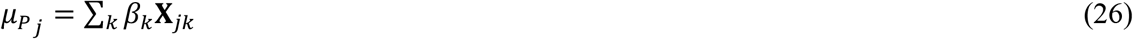

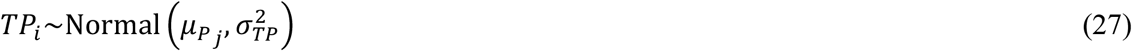

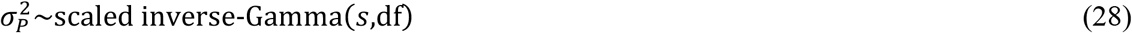

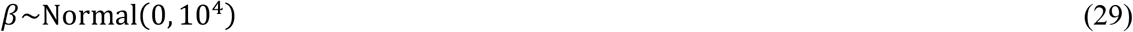

where *μ*_*j*_ is the expected value of the *j*th observation, **X** is a model matrix including an intercept and the two explanatory variables (NPP [log-transformed] and growing season length) and an associated vector of parameters *β*. We used an uninformative normal prior with a mean of 0 and a variance of 10^4^ for *β* and a weakly informative-scaled inverse gamma prior with scale *s* = 1 and two degrees of freedom (d.f. = 2) for the residual variance.

### Model implementation and assessment of model fit

We used posterior predictive checks to assess the global fit of the models to the data (69). As a measure of global model fit, we computed a posterior predictive *P* value (PPP) that compares the residual sum of squares (RSS) based on the observed data to RSS generated from the posterior predictive distribution of the model. Values of PPP close to 0.5 indicate that the model fits the observed data, whereas values close to 0 or 1 indicate the opposite. We report the marginal variance, *r*_*m*_^2^, i.e., explained by the fixed factors, as well as the conditional variance, *r*_*c*_^2^, i.e., explained by the fixed and random factors combined (70). Finally, we provide the median of all model estimates along with 50 and 90% Equal-Tailed Intervals (ETIs) as measures of uncertainty (71). As a measure of support for the effects of the explanatory variables, we calculated the fraction of posterior samples with the same sign as the median (*p*_*d*_). This metric quantifies the probability that the effect of a given explanatory variable differs from zero.

For both models we provide partial residual plots to illustrate the relationships of trophic position with NPP and growing season length and for visual inspection of model fit to single observations. To enhance the interpretation of the partial residual plots, we standardized the two predictor variables to zero mean and unit variance so that the effect of each predictor variable is shown while conditioning on the mean value of other predictor variable.

All analyses were conducted in *R* [version 4.0.3 (58)]. The Bayesian models were implemented in *JAGS* (version 4.3) and run in *R* through the *rjags* package [version 4-10 (72)]. We ran five parallel chains for the models. Each chain was run for 201,000 iterations with an adaptive burn-in phase of 1,000 iterations and a thinning interval of 100 iterations, resulting in 2,000 samples per chain, corresponding to 10,000 samples from the posterior distribution. Initial values were drawn randomly from the prior distributions using package *LaplacesDemon* [version 16.1.6 (73)]. The chains were checked for convergence, temporal autocorrelation and effective sample size using the *coda* package [version 0.19-4 (74)]. The residuals were checked for normality and variance homogeneity. We assessed potential collinearity among predictors using variance inflation factors (75).

## Acknowledgements

This study was funded by the Norway grants under the Polish-Norwegian Research Program administered by the National Research Centre for Research and Development in Poland (project GLOBE No POL-NOR/198352/85/2013). J.A. was funded by the German Academic Exchange Service in the framework of a post doctorate fellowship grant (DAAD, No 91568794).

## Author contributions

J.A. and N.S. conceived the study. J.A. compiled data from the literature. J.A., A.S., A.M., E.I., L.D. collected material and took samples from museum collections for stable isotope analyses. H.B., D.G.D. and K.A.H. conducted stable isotope analysis. J.A. developed the analytical tools, processed and analyzed the data and wrote the initial draft of the manuscript with input from N.S., H.B., K.A.H. and S.A.F.; J.A., H.B., K.A.H., D.G.D., A.S., J.S., A.Z., A.M., E.I., L.D., R.K., G.L., D.M., J.W., T.Z.-K., S.A.F., E.R. and N.S. reviewed and edited subsequent versions of the manuscript.

## References

1. J. M. Tylianakis, R. K. Didham, J. Bascompte, D. A. Wardle, Global change and species interactions in terrestrial ecosystems. Ecol. Lett. 11, 1351–1363 (2008).

2. J. A. Estes, et al., Trophic downgrading of planet Earth. Science 333, 301–306 (2011).

3. E. G. Ritchie, et al., Ecosystem restoration with teeth: what role for predators? Trends Ecol. Evol. 27, 265–271 (2012).

4. T. J. Bartley, et al., Food web rewiring in a changing world. Nat. Ecol. Evol. 3, 345–354 (2019).

5. M. K. Gutgesell, et al., On the dynamic nature of omnivory in a changing world. BioScience 72, 416–430 (2022).

6. S. Wang, U. Brose, D. Gravel, Intraguild predation enhances biodiversity and functioning in complex food webs. Ecology 100, e02616 (2019).

7. T. Kastner, et al., Land use intensification increasingly drives the spatiotemporal patterns of the global human appropriation of net primary production in the last century. Glob. Change Biol. 28, 307–322 (2022).

8. D. S. LeBauer, K. K. Treseder, Nitrogen limitation of net primary productivity in terrestrial ecosystems is globally distributed. Ecology 89, 371–379 (2008).

9. T. M. Newsome, et al., The ecological effects of providing resource subsidies to predators. Glob. Ecol. Biogeogr. 24, 1–11 (2015).

10. H. Liu, C. Lu, S. Wang, F. Ren, H. Wang, Climate warming extends growing season but not reproductive phase of terrestrial plants. Glob. Ecol. Biogeogr. 30, 950–960 (2021).

11. D. M. Post, G. Takimoto, Proximate structural mechanisms for variation in food-chain length. Oikos 116, 775–782 (2007).

12. C. L. Ward, K. S. McCann, A mechanistic theory for aquatic food chain length. Nat. Commun. 8, 2028 (2017).

13. S. Diehl, M. Feissel, Intraguild prey suffer from enrichment of their resources: A microcosm experiment with ciliates. Ecology 82, 2977–2983 (2001).

14. A. Sentis, J.-L. Hemptinne, J. Brodeur, Towards a mechanistic understanding of temperature and enrichment effects on species interaction strength, omnivory and food-web structure. Ecol. Lett. 17, 785–793 (2014).

15. J. L. Blois, P. L. Zarnetske, M. C. Fitzpatrick, S. Finnegan, Climate change and the past, present, and future of biotic interactions. Science 341, 499–504 (2013).

16. D. A. Fordham, et al., Using paleo-archives to safeguard biodiversity under climate change. Science 369, eabc5654 (2020).

17. T. Sacco, B. Van Valkenburgh, Ecomorphological indicators of feeding behaviour in the bears (Carnivora: Ursidae). J. Zool. 263, 41–54 (2004).

18. C. T. Robbins, et al., Ursids evolved early and continuously to be low-protein macronutrient omnivores. Sci. Rep. 12, 15251 (2022).

19. J. Berger, P. B. Stacey, L. Bellis, M. P. Johnson, A mammalian predator-prey imbalance: grizzly bear and wolf extinction affect avian neotropical migrants. Ecol. Appl. 11, 947–960 (2001).

20. J. B. Grinath, B. D. Inouye, N. Underwood, Bears benefit plants via a cascade with both antagonistic and mutualistic interactions. Ecol. Lett. 18, 164–173 (2015).

21. A. García-Rodríguez, et al., The role of the brown bear Ursus arctos as a legitimate megafaunal seed disperser. Sci. Rep. 11, 1282 (2021).

22. O. J. Schmitz, D. Hawlena, G. C. Trussell, Predator control of ecosystem nutrient dynamics. Ecol. Lett. 13, 1199–1209 (2010).

23. W. W. Deacy, et al., Phenological synchronization disrupts trophic interactions between Kodiak brown bears and salmon. Proc. Natl. Acad. Sci. 114, 10432–10437 (2017).

24. J. Albrecht, et al., Data and Code from “Trophic adaptation of large terrestrial omnivores to global change.” figshare 10.6084/m9.figshare.25671048 (2024).

25. K. Hobson, H. Welch, Determination of trophic relationships within a high Arctic marine food web using δ13C and δ15N analysis. Mar. Ecol. Prog. Ser. 84, 9–18 (1992).

26. K. A. Hobson, I. Stirling, Low variation in blood δ13C among Hudson bay polar bears: Implications for metabolism and tracing terrestrial foraging. Mar. Mammal Sci. 13, 359–367 (1997).

27. M. A. Ramsay, K. A. Hobson, Polar Bears Make Little Use of Terrestrial Food Webs: Evidence from Stable-Carbon Isotope Analysis. (2008).

28. K. A. Hobson, I. Stirling, D. S. Andriashek, Isotopic homogeneity of breath CO 2 from fasting and berry-eating polar bears: implications for tracing reliance on terrestrial foods in a changing Arctic. Can. J. Zool. 87, 50–55 (2009).

29. C. Körner, P. Möhl, E. Hiltbrunner, Four ways to define the growing season. Ecol. Lett. ele.14260 (2023). 10.1111/ele.14260.

30. F.-E. Wielgolaski, Starting dates and basic temperatures in phenological observations of plants. Int. J. Biometeorol. 42, 158–168 (1999).

31. M. M. Delgado, et al., The seasonal sensitivity of brown bear denning phenology in response to climatic variability. Front. Zool. 15, 41 (2018).

32. N. L. Fowler, J. L. Belant, G. Wang, B. D. Leopold, Ecological plasticity of denning chronology by American black bears and brown bears. Glob. Ecol. Conserv. 20, e00750 (2019).

33. J. L. Belant, K. Kielland, E. H. Follmann, L. G. Adams, Interspecific resource partitioning in sympatric Ursids. Ecol. Appl. 16, 2333–2343 (2006).

34. H. Bocherens, Isotopic tracking of large carnivore palaeoecology in the mammoth steppe. Quat. Sci. Rev. 117, 42–71 (2015).

35. T. T. Männel, K. Auerswald, H. Schnyder, Altitudinal gradients of grassland carbon and nitrogen isotope composition are recorded in the hair of grazers. Glob. Ecol. Biogeogr. 16, 583–592 (2007).

36. H. Bocherens, D. Drucker, Trophic level isotopic enrichment of carbon and nitrogen in bone collagen: case studies from recent and ancient terrestrial ecosystems. Int. J. Osteoarchaeol. 13, 46–53 (2003).

37. M. Krapp, R. M. Beyer, S. L. Edmundson, P. J. Valdes, A. Manica, A statistics-based reconstruction of high-resolution global terrestrial climate for the last 800,000 years. Sci. Data 8, 228 (2021).

38. C. A. Welch, J. Keay, K. C. Kendall, C. T. Robbins, Constraints on frugivory by bears. Ecology 78, 1105–1119 (1997).

39. J. Matsubayashi, et al., Major decline in marine and terrestrial animal consumption by brown bears (Ursus arctos). Sci. Rep. 5, 9203 (2015).

40. D. C. Rinker, N. K. Specian, S. Zhao, J. G. Gibbons, Polar bear evolution is marked by rapid changes in gene copy number in response to dietary shift. Proc. Natl. Acad. Sci. 116, 13446–13451 (2019).

41. M. J. de Jong, et al., Range-wide whole-genome resequencing of the brown bear reveals drivers of intraspecies divergence. Commun. Biol. 6, 153 (2023).

42. M. Baca, et al., Retreat and extinction of the Late Pleistocene cave bear (Ursus spelaeus sensu lato). Sci. Nat. 103, 92 (2016).

43. Ö. E. Can, N. D’Cruze, D. L. Garshelis, J. Beecham, D. W. Macdonald, Resolving human-bear conflict: A global survey of countries, experts, and key factors. Conserv. Lett. 7, 501–513 (2014).

44. K. Bojarska, N. Selva, Spatial patterns in brown bear Ursus arctos diet: the role of geographical and environmental factors. Mammal Rev. 42, 120–143 (2012).

45. M. Krofel, M. Špacapan, K. Jerina, Winter sleep with room service: denning behaviour of brown bears with access to anthropogenic food. J. Zool. 302, 8–14 (2017).

46. A. J. Mikkelsen, et al., Testing foraging optimization models in brown bears: Time for a paradigm shift in nutritional ecology? Ecology e4228 (2023). 10.1002/ecy.4228.

47. A. M. Pagano, “Polar Bear Foraging Behavior” in Ethology and Behavioral Ecology of Sea Otters and Polar Bears, Ethology and Behavioral Ecology of Marine Mammals., R. W. Davis, A. M. Pagano, Eds. (Springer International Publishing, 2021), pp. 247–267.

48. Y. Malhi, et al., Megafauna and ecosystem function from the Pleistocene to the Anthropocene. Proc. Natl. Acad. Sci. 113, 838–846 (2016).

49. W. D. Kissling, et al., Establishing macroecological trait datasets: digitalization, extrapolation, and validation of diet preferences in terrestrial mammals worldwide. Ecol. Evol. 4, 2913–2930 (2014).

50. S. Running, M. Zhao, MOD17A3HGF MODIS/Terra Net Primary Production Gap-Filled Yearly L4 Global 500 m SIN Grid V006. NASA EOSDIS Land Processes DAAC. 10.5067/MODIS/MOD17A3HGF.006. Deposited 2019.

51. D. N. Karger, et al., Climatologies at high resolution for the earth’s land surface areas. Sci. Data 4, 170122 (2017).

52. S. Xu, A property of geometric mean regression. Am. Stat. 68, 277–281 (2014).

53. A. Huang, M. P. Wand, Simple marginally noninformative prior distributions for covariance matrices. Bayesian Anal. 8, 439–452 (2013).

54. A. Gelman, Prior distributions for variance parameters in hierarchical models (comment on article by Browne and Draper). Bayesian Anal. 1, 515–533 (2006).

55. J. C. Douma, J. T. Weedon, Analysing continuous proportions in ecology and evolution: A practical introduction to beta and Dirichlet regression. Methods Ecol. Evol. 10, 1412–1430 (2019).

56. A. D. Barnosky, E. L. Lindsey, Timing of Quaternary megafaunal extinction in South America in relation to human arrival and climate change. Quat. Int. 217, 10–29 (2010).

57. J. Haslett, A. Parnell, A Simple Monotone Process with Application to Radiocarbon-Dated Depth Chronologies. J. R. Stat. Soc. Ser. C Appl. Stat. 57, 399–418 (2008).

58. R Development Core Team, R: A language and environment for statistical computing (R Foundation for Statistical Computing, 2020).

59. P. J. Reimer, et al., IntCal13 and Marine13 Radiocarbon Age Calibration Curves 0–50,000 Years cal BP. Radiocarbon 55, 1869–1887 (2013).

60. R. Longin, New method of collagen extraction for radiocarbon dating. Nature 230, 241–242 (1971).

61. H. Bocherens, et al., Paleobiological implications of the isotopic signatures (13C, 15N) of fossil mammal collagen in Scladina cave (Sclayn, Belgium). Quat. Res. 48, 370–380 (1997).

62. M. J. DeNiro, Postmortem preservation and alteration of in vivo bone collagen isotope ratios in relation to palaeodietary reconstruction. Nature 317, 806–809 (1985).

63. S. H. Ambrose, Preparation and characterization of bone and tooth collagen for isotopic analysis. J. Archaeol. Sci. 17, 431–451 (1990).

64. M. Leonardi, E. Y. Hallett, R. Beyer, M. Krapp, A. Manica, pastclim 1.2: an R package to easily access and use paleoclimatic reconstructions. Ecography 2023, e06481 (2023).

65. Earth Resources Observation and Science (EROS) Center, Global Multi-resolution Terrain Elevation Data 2010 (GMTED2010). U.S. Geological Survey. 10.5066/F7J38R2N. Deposited 2017.

66. S. O. Rasmussen, et al., A stratigraphic framework for abrupt climatic changes during the Last Glacial period based on three synchronized Greenland ice-core records: refining and extending the INTIMATE event stratigraphy. Quat. Sci. Rev. 106, 14–28 (2014).

67. M. Walker, et al., Formal ratification of the subdivision of the Holocene Series/Epoch (Quaternary System/Period): two new Global Boundary Stratotype Sections and Points (GSSPs) and three new stages/subseries. Episodes 41, 213–223 (2018).

68. D. M. Post, Using stable isotopes to estimate trophic position: models, methods, and assumptions. Ecology 83, 703–718 (2002).

69. A. Gelman, et al., Bayesian data analysis, 3rd Ed. (Chapman and Hall/CRC, 2013).

70. S. Nakagawa, P. C. D. Johnson, H. Schielzeth, The coefficient of determination R2 and intra-class correlation coefficient from generalized linear mixed-effects models revisited and expanded. J. R. Soc. Interface 14, 20170213 (2017).

71. R. McElreath, Statistical rethinking: a Bayesian course with examples in R and Stan, Second edition (CRC Press, 2020).

72. M. Plummer, rjags: Bayesian graphical models using MCMC (2022).

73. Statisticat, LLC., LaplacesDemon: Complete environment for Bayesian inference (Bayesian-Inference.com, 2021).

74. M. Plummer, N. Best, K. Cowles, K. Vines, CODA: convergence diagnosis and output analysis for MCMC. R News 6, 7–11 (2006).

75. C. F. Dormann, et al., Collinearity: a review of methods to deal with it and a simulation study evaluating their performance. Ecography 36, 27–46 (2013).

76. S. Kumar, et al., TimeTree 5: An Expanded Resource for Species Divergence Times. Mol. Biol. Evol. 39, msac174 (2022).

