## Supplementary Information for "Trophic adaptation of large terrestrial omnivores to global change"

**Table S1. Summary of dietary classification scheme and associated correction factors.**

| diet1 | diet2 | diet3 | $c_D$ | $c_E$ | comment | $c_D$ reference | $c_E$ reference |
|---|---|---|---|---|---|---|---|
| animals | vertebrates | endotherm_vertebrates | 3.125 | 14.200 | – | (1–4) | (2) |
| animals | vertebrates | ectotherm_vertebrates | 3.125 | 14.200 | – | (1–4) | (2) |
| animals | vertebrates | fish_vertebrates | 40.800 | 14.200 | – | – | – |
| animals | vertebrates | unknown_vertebrates | 3.125 | 14.200 | mean of endo and ecto vertebrates | – | – |
| animals | invertebrates | aquatic_invertebrates | 1.100 | 14.500 | assumed same as terrestrial invertebrates | (1, 3, 4) | (3, 4) |
| animals | invertebrates | terrestrial_invertebrates | 1.100 | 14.500 | – | (1, 3, 4) | (3, 4) |
| animals | invertebrates | unknown_invertebrates | 1.100 | 14.500 | mean of aquatic and terrestrial invertebrates | – | – |
| animals | unknown_animal_material | unknown_animal_material | 2.113 | 14.350 | mean of vertebrates and invertebrates | – | – |
| plants | reproductive_plant_material | fruit | 1.064 | 18.100 | – | (1, 5) | (6) |
| plants | reproductive_plant_material | seeds | 1.450 | 26.700 | – | (1, 5) | (6) |
| plants | reproductive_plant_material | flowers_nectar_pollen | 0.270 | 8.400 | assumed same as forbs, herbs, legumes | (1) | (3, 4) |
| plants | reproductive_plant_material | unknown_reproductive_plant_material | 0.928 | 17.733 | mean of reproductive plant materials | – | – |
| plants | vegetative_plant_material | forbs_herbs_legumes | 0.270 | 8.400 | – | (1) | (3, 4) |
| plants | vegetative_plant_material | grass | 0.230 | 6.300 | – | (1) | (3, 4) |
| plants | vegetative_plant_material | roots_tubers_bulbs | 0.800 | 16.500 | – | (1) | (6) |
| plants | vegetative_plant_material | leaves_branches_bark | 0.270 | 8.400 | assumed same as forbs, herbs, legumes | (1) | (3, 4) |
| plants | vegetative_plant_material | unknown_vegetative_plant_material | 0.393 | 9.900 | mean of vegetative plant materials | – | – |
| plants | unknown_plant_material | unknown_plant_material | 0.660 | 13.817 | mean of reproductive and vegetative plant materials | – | – |
| fungi_lichens_bryophytes_algae | fungi_lichens_bryophytes_algae | fungi_lichens_bryophytes_algae | 0.270 | 8.400 | assumed same as forbs, herbs, legumes | (1) | (3, 4) |
| other | other | other | 0.000 | 0.000 | – | – | – |

Note: Diet1, Diet2, and Diet3 are dietary classifications at different resolutions. Provided are also correction factors (and literature sources) that account for differences in digestibility ($c_D$) and energy content ($c_E$) of food items in Diet3.

**Table S2. Summary of Bayesian hierarchical model testing for relationships of the trophic position (i.e., relative dietary energy contribution of animal prey) with net primary productivity (NPP), growing season length (GSL), and competition with sympatric bear species across the geographic ranges of the seven extant terrestrial bear species.**

| Source of variance | Median | 5% | 95% | $p_d$ | PSRF | $N_{eff}$ |
|---|---|---|---|---|---|---|
| (**a**) Sub-model I ($V \sim F$), PPP = 0.4986 | | | | | | |
| Intercept | 0.428 | 0.291 | 0.56 | 1 | 1 | 10000 |
| Slope | 1.32 | 1.23 | 1.4 | 1 | 1 | 10000 |
| $\sigma^2_{intercept}$ | 0.784 | 0.632 | 0.99 | – | 1 | 10292 |
| $\sigma^2_{slope}$ | 0.282 | 0.226 | 0.355 | – | 1 | 10319 |
| $\rho_{intercept,slope}$ | 0.955 | 0.938 | 0.967 | – | 1 | 10000 |
| $\sigma^2_{residual}$ | 0.723 | 0.666 | 0.785 | – | 1 | 10000 |
| $r^2$ | 0.828 | 0.809 | 0.848 | – | 1 | 9457 |
| (**b**) Sub-model II (Intercept only), PPP = 0.5073 | | | | | | |
| Intercept | -0.787 | -1.15 | -0.459 | 0.999 | 1 | 10000 |
| $\sigma^2_{species}$ | 0.155 | 0.0402 | 0.613 | – | 1.01 | 7522 |
| $\sigma^2_{study}$ | 0.518 | 0.397 | 0.667 | – | 1 | 10000 |
| $\sigma^2_{residual}$ | 0.134 | 0.0974 | 0.19 | – | 1 | 10000 |
| $r^2_{marginal}$ | 3.88E-30 | 1.84E-32 | 2.17E-29 | – | 1 | 0 |
| $r^2_{conditional}$ | 0.633 | 0.39 | 0.775 | – | 1 | 9435 |
| (**c**) Sub-model II (GSL + NPP + competition), PPP = 0.5127 | | | | | | |
| Intercept | -0.549 | -0.941 | -0.165 | 0.982 | 1 | 10000 |
| Growing season length (GSL) | -0.414 | -0.574 | -0.259 | 1 | 1 | 10017 |
| Net primary productivity (NPP) | -0.183 | -0.305 | -0.0638 | 0.993 | 1 | 10795 |
| Competition (subordinate) | -0.501 | -0.836 | -0.155 | 0.992 | 1 | 10000 |
| Competition (dominant | 0.0534 | -0.256 | 0.373 | 0.612 | 1 | 10000 |
| $\sigma^2_{species}$ | 0.211 | 0.0196 | 0.969 | – | 1 | 10000 |
| $\sigma^2_{study}$ | 0.319 | 0.224 | 0.432 | – | 1 | 10000 |
| $\sigma^2_{residual}$ | 0.137 | 0.0991 | 0.194 | – | 1 | 10000 |
| $r^2_{marginal}$ | 0.275 | 0.145 | 0.394 | – | 1 | 10000 |
| $r^2_{conditional}$ | 0.626 | 0.336 | 0.805 | – | 1 | 10000 |
| (**d**) Sub-model II (Two-component model), PPP = 0.5094 | | | | | | |
| Intercept | -0.736 | -1.41 | -0.257 | 0.983 | 1 | 10000 |
| $GSL_{ij}$ – mean$(GSL_{ij})_j$ | -0.441 | -0.596 | -0.284 | 1 | 1 | 10000 |
| mean$(GSL_{ij})_j$ | 0.0464 | -0.803 | 1.06 | 0.542 | 1 | 10000 |
| $NPP_{ij}$ – mean$(NPP_{ij})_j$ | -0.178 | -0.297 | -0.0574 | 0.991 | 1 | 9240 |
| mean$(NPP_{ij})_j$ | -0.234 | -1.08 | 0.76 | 0.69 | 1 | 10000 |
| Competition (subordinate) | -0.526 | -0.867 | -0.18 | 0.994 | 1 | 10000 |
| Competition (dominant) | 0.0675 | -0.238 | 0.378 | 0.641 | 1 | 10000 |
| $\sigma^2_{species}$ | 0.281 | 0.0397 | 1.4 | – | 1.02 | 9626 |
| $\sigma^2_{study}$ | 0.312 | 0.22 | 0.42 | – | 1 | 10680 |
| $\sigma^2_{residual}$ | 0.138 | 0.1 | 0.195 | – | 1 | 10719 |
| $r^2_{marginal}$ | 0.252 | 0.129 | 0.396 | – | 1 | 10000 |
| $r^2_{conditional}$ | 0.575 | 0.311 | 0.768 | – | 1 | 10000 |

Given are estimated model parameters [median and 90% Equal-Tailed Intervals (ETIs)], and convergence diagnostics [potential scale reduction factor (PSRF) and effective sample size ($N_{eff}$)] for all model parameters. The fraction of posterior samples with the same sign as the median ($p_d$) is provided as a measure of support for the effects of the explanatory variables. PPP-values for each sub-model indicate how well model simulations fit the observed data. Values of PPP close to 0.5 indicate that the model fits the observed data, while values close to 0 or 1 indicate the opposite.

**Table S3. Summary of Bayesian hierarchical model testing for relationships of trophic position (i.e., trophic level) with net primary productivity (NPP) and growing season length (GSL) in the brown bear (*Ursus arctos*) from the Late Pleistocene to the Holocene.**

| Source of variance | Median | 5% | 95% | $p_d$ | PSRF | $N_{eff}$ |
|---|---|---|---|---|---|---|
| (**a**) Sub-model I | | | | | | |
| Effect of elevation, PPP = 0.5063 | | | | | | |
| Tissue type | | | | | | |
| Vegetation | 2.87 | 2.47 | 3.27 | 1 | 1 | 10000 |
| Sheep wool | 6.46 | 5.97 | 6.94 | 1 | 1 | 10459 |
| Goat hair | 5.93 | 5.4 | 6.47 | 1 | 1 | 10000 |
| Cattle hair | 6.09 | 5.31 | 6.85 | 1 | 1 | 10000 |
| Elevation ($\beta_{\text{Elevation}}$) | -0.0013 | -0.00153 | -0.00106 | 1 | 1 | 10000 |
| $\sigma^2_{\text{residual}}$ | 1.15 | 0.87 | 1.55 | – | 1 | 10000 |
| Effect of sampled material (bone vs. tooth), PPP = 0.4976 | | | | | | |
| $\mu_T$ | 1.51 | 1.31 | 1.71 | 1 | 1 | 10000 |
| $\sigma^2_T$ | 0.542 | 0.373 | 0.84 | – | 1 | 10988 |
| Trophic discrimination factor, PPP = 0.5102 | | | | | | |
| $\mu_\Delta$ | 3.9 | 3.33 | 4.54 | 1 | 1 | 9157 |
| $\sigma^2_\Delta$ | 0.839 | 0.353 | 1.67 | – | 1 | 9279 |
| (**b**) Sub-model II, PPP = 0.4381 | | | | | | |
| Intercept | 2.34 | 2.22 | 2.49 | 1 | 1 | 10765 |
| Growing season length (GSL) | -0.176 | -0.308 | -0.0317 | 0.969 | 1 | 10000 |
| Net primary productivity (NPP) | -0.183 | -0.323 | -0.0352 | 0.972 | 1 | 10000 |
| $\sigma^2_{\text{residual}}$ | 0.0187 | 0.000443 | 0.15 | – | 1 | 10000 |
| $r^2$ | 0.78 | 0.226 | 0.994 | – | 1 | 9652 |

Given are estimated model parameters [median and 90% Equal-Tailed Intervals (ETIs)], and convergence diagnostics [potential scale reduction factor (PSRF) and effective sample size ($N_{eff}$)] for all model parameters. The fraction of posterior samples with the same sign as the median ($p_d$) is provided as a measure of support for the effects of the explanatory variables. PPP-values for each sub-model indicate how well model simulations fit the observed data. Values of PPP close to 0.5 indicate that the model fits the observed data, while values close to 0 or 1 indicate the opposite.

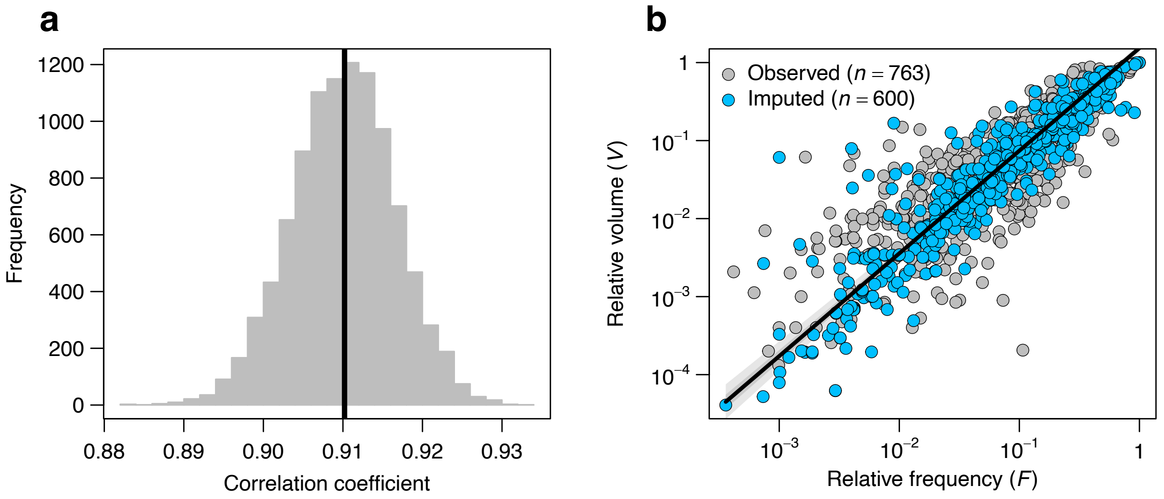


**Figure S1. Relationship between relative frequency (*F*) and relative volume (*V*) of dietary food items. a**, Median (black line) and posterior distribution of estimated correlation coefficient between *F* and *V* based on sub-model I in the macroecological analysis (Table S2a). **b**, Geometric mean regression relationship between *F* and *V* for observed (white dots) and imputed (blue dots) data. Black line, dark and light gray bands represent the estimated relationship and uncertainty (median, 50% and 90% ETIs, respectively).

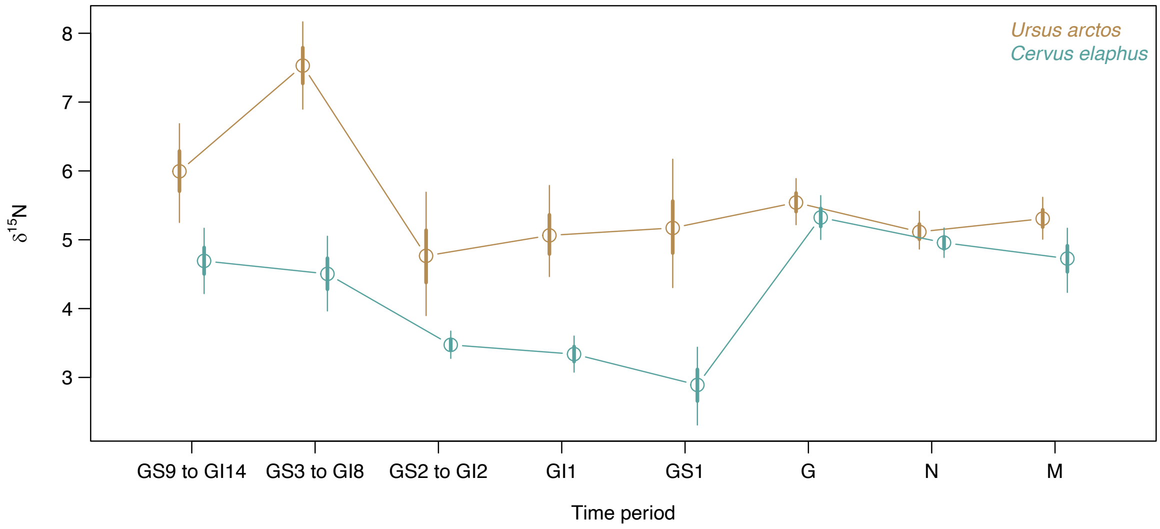


**Supplementary Figure 2. Estimated bias-corrected $\delta^{15}N$ value of red deer and brown bear in each of the eight time periods.** Median (circles) and associated 50%, and 90% Equal-Tailed Intervals (thick and thin lines) of the bias-corrected $\delta^{15}N$ values of red deer (purple) and brown bear (orange) based on sub-model I in the paleoecological model (Table S3a).

### References

1. D. G. Hewitt, C. T. Robbins, Estimating Grizzly Bear Food Habits from Fecal Analysis. *Wildl. Soc. Bull.* **24**, 547–550 (1996).
2. T. Johansen, “The diet of the brown bear (Ursus arctos) in central Sweden,” Norwegian University of Science and Technology, Trondheim (1997).
3. B. Dahle, O. Sørensen, E. Wedul, J. E. Swenson, F. Sandegren, The diet of brown bears Ursus arctos in central Scandinavia: effect of access to free-ranging domestic sheep Ovis aries. *Wildl. Biol.* **4**, 147–158 (1998).
4. I.-L. Persson, S. Wikan, J. E. Swenson, I. Mysterud, The diet of the brown bear Ursus arctos in the Pasvik Valley, northeastern Norway. *Wildl. Biol.* **7**, 27–37 (2001).
5. K. Bojarska, N. Selva, Correction factors for important brown bear foods in Europe. *Ursus* **24**, 13–15 (2013).
6. G. T. Pritchard, C. T. Robbins, Digestive and metabolic efficiencies of grizzly and black bears. *Can. J. Zool.* **68**, 1645–1651 (1990).
